# Cross-Regional Trajectory Analysis Reveals Distinct Temporal Progression of Shared Proteomic Programs in Alzheimer’s Disease

**DOI:** 10.64898/2026.08.20.746024

**Authors:** Tianchuan Gao, Yurika Upadhyaya, Yanling Pan, Kwangsik Nho, Andrew J. Saykin, Jingwen Yan

## Abstract

Alzheimer’s pathology progresses through anatomically distinct brain regions, yet how molecular programs are temporally organized across vulnerable regions remains poorly understood. Here, we reconstructed continuous proteomic trajectories from matched dorsolateral prefrontal cortex (DLPFC) and superior temporal gyrus (STG) proteomes to investigate cross-regional molecular progression in AD. The inferred trajectories closely recapitulated established neuropathological staging while remaining independent of age, sex, and race. Across regions, a mitochondrial bioenergetic protein subset declined substantially earlier in the STG than in the DLPFC, revealing a previously unrecognized regional temporal offset in metabolic dysfunction. In parallel, a broadly shared proteomic program progressed in both regions but more rapidly in the STG, indicating that shared molecular responses differ in their temporal progression despite overall coordination. Together, these findings demonstrate that regional vulnerability in AD is reflected not only by the molecular programs involved but also by their temporal organization during disease progression.

## Introduction

Alzheimer’s disease (AD) is characterized by selective vulnerability of anatomically distinct brain regions. Neuropathological studies have shown that amyloid-β deposition, tau pathology, and neurodegeneration involve different cortical regions in an ordered sequence during disease progression^1,2^. Although this regional organization has been extensively characterized neuropathologically, the molecular mechanisms underlying selective regional vulnerability remain incompletely understood^3^. Elucidating these molecular mechanisms is essential for understanding the biological processes governing disease progression across the brain. The rapid expansion of large-scale human brain proteomic datasets has enabled systematic characterization of disease-associated molecular alterations across multiple vulnerable cortical regions^4–6^, providing an opportunity to investigate how molecular responses are organized across the AD brain.

Recent cross-regional proteomic studies have progressively advanced our understanding of this molecular organization. Differential protein abundance analyses first demonstrated that many AD-associated protein alterations are shared across vulnerable cortical regions despite substantial differences in regional neuropathological burden^7^. The magnitude of these protein abundance changes, however, often differed between regions, indicating that common disease-associated alterations are not uniformly manifested across the cortex. Another analyses with matched dorsolateral prefrontal cortex (DLPFC) and superior temporal gyrus (STG) samples demonstrated that these shared disease-associated protein abundance changes were not only shared but also highly correlated between cortical regions, providing direct evidence for coordinated molecular responses across vulnerable cortical regions^6^. Going beyond individual proteins, protein co-expression analyses further extended these observations and demonstrated that disease-associated proteins are organized into functional molecular modules representing major biological processes, including synaptic function, mitochondrial metabolism, immune activation, and proteostasis^5,8^. Subsequent cross-regional network analyses further demonstrated that many of these molecular programs are shared across cortical regions, indicating that coordinated disease-associated responses are organized into common higher-order biological systems rather than isolated protein alterations^8^. The reproducibility of these shared molecular programs across independent cohorts further established them as robust features of AD proteomic remodeling^5^.

Despite these advances, the temporal organization of cross-regional proteomic progression remains largely unknown. Existing cross-regional proteomic studies primarily compared protein abundance or co-expression patterns across diagnostic groups or neuropathological stages, providing cross-sectional descriptions of disease-associated molecular alterations. While these studies established the molecular programs shared across vulnerable cortical regions, they did not determine whether these programs emerge synchronously or instead exhibit distinct regional temporal progression during AD. To address this question, we investigated the temporal progression of proteomic programs across vulnerable cortical regions using matched dorsolateral prefrontal cortex (DLPFC) and superior temporal gyrus (STG) proteomes from the AMP-AD Diverse Cohorts resource^6^. By reconstructing a continuous molecular progression axis, we compared the temporal trajectories of shared proteomic programs between regions to determine whether they followed similar or distinct patterns of progression. This framework provides a temporal perspective on regional proteomic progression during AD.

## Results

### Study design and cohort characteristics

We assembled a cross-regional proteomic cohort of 215 donors with matched dorsolateral prefrontal cortex (DLPFC) and superior temporal gyrus (STG) tandem mass tag (TMT) proteomic profiles from the AMP-AD Diverse Cohorts study (Fig. 1A–C). Of the 579 DLPFC and 242 STG donors in the complete dataset, 215 had matched proteomic measurements from both brain regions. The remaining donors comprised a non-overlapping DLPFC-only cohort (n = 364) and a small STG-only cohort (n = 27) (Fig. 1B). The DLPFC-only cohort was reserved for independent validation of trajectory reproducibility.

**Fig. 1.**
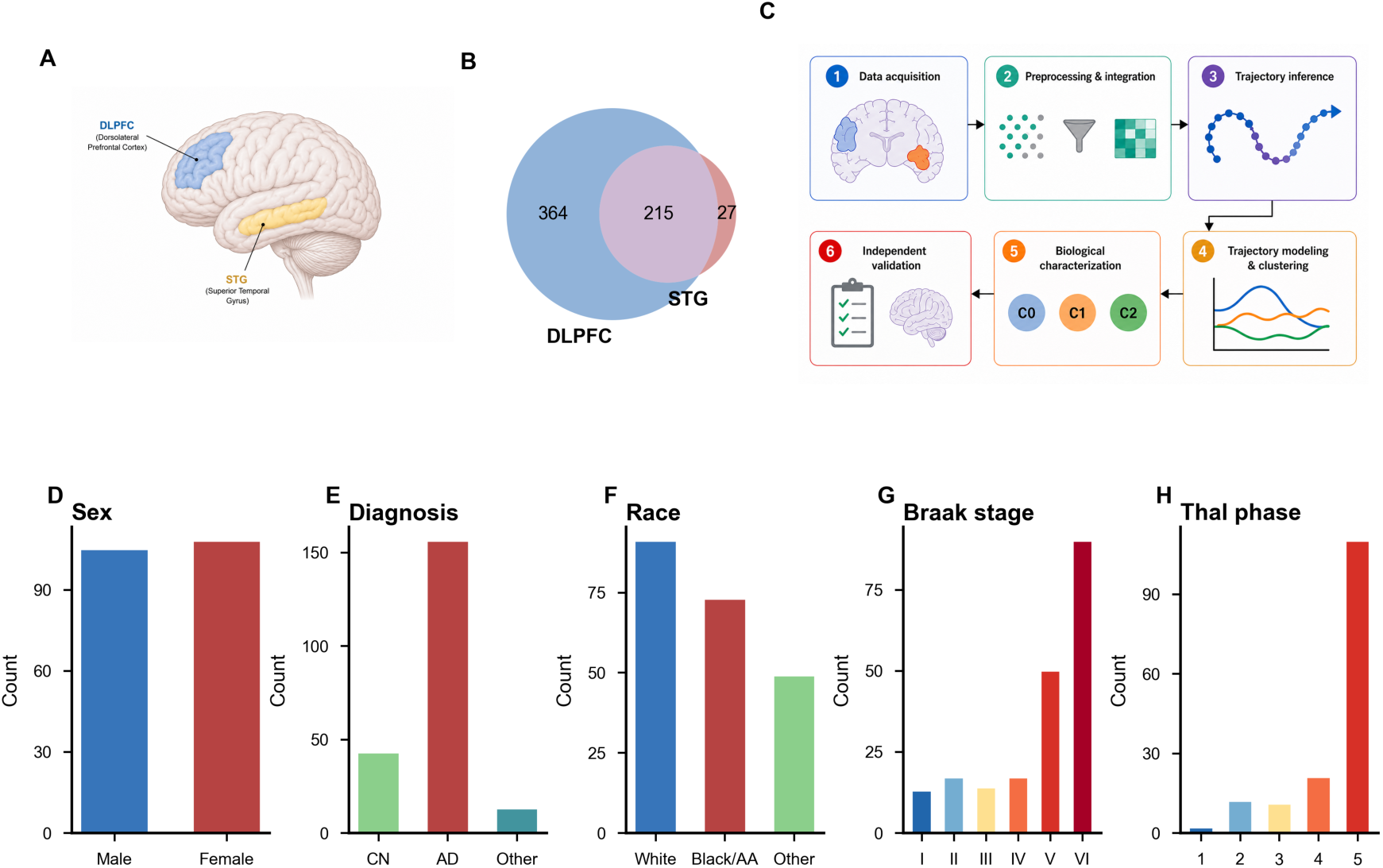
Study design and cohort characteristics. (A) Two profiled cortical regions: dorsolateral prefrontal cortex (DLPFC) and superior temporal gyrus (STG). (B) Donor overlaps between two brain regions. (C) Analytical workflow for cross-regional proteomic trajectory analysis and independent validation. (D-F) Sex, diagnosis, and race/ethnicity distributions among donors shared across brain regions. (G–H) Braak stage and Thal phase distributions among donors shared across brain regions.

Following phenotype and quality-control filtering, 213 of the 215 matched donors were retained for trajectory analysis. Two donors were excluded because of missing phenotype annotations or failure to meet quality-control criteria. The final cohort included 105 males and 108 females, with 91 White, 73 Black or African American, and 49 individuals from other racial or ethnic backgrounds. Diagnostic groups included 156 Alzheimer’s disease cases, 43 cognitively normal controls, and 14 donors with other or unknown diagnoses (Fig. 1D–F; Supplementary Fig. 1). Braak neurofibrillary tangle stage and Thal amyloid phase spanned the full spectrum of neuropathological severity represented in the cohort (Fig. 1G,H). In total, the matched cross-regional dataset comprised 18,036 region-specific protein features, corresponding to 9,018 unique protein groups measured across the two brain regions. Detailed cohort characteristics, missingness summaries, and inclusion criteria are provided in Supplementary Table 1.

The matched DLPFC–STG cohort enabled direct comparison of proteomic progression between two cortical regions, whereas the independent DLPFC-only cohort provided an opportunity to evaluate the reproducibility of the identified trajectory patterns. The overall study design and analytical workflow are summarized in Fig. 1C.

### Proteomic pseudotime recapitulates Alzheimer’s disease progression

PHATE embedding of the cross-regional proteome, after removing the effects of age at death, sex, and PMI, revealed a prominent low-dimensional trajectory structure across the 213 matched donors (Fig. 2A). Pseudotime was assigned to each donor based on their relative position on the trajectory curve. This trajectory shows a continuous axis from 0 (low-pathology, control-enriched) to 1 (high-pathology, AD-enriched). AD individuals occupied significantly higher pseudotime positions than cognitively normal controls (Mann-Whitney U *p* = 3.2 × 10^−1^^2^, Fig. 2B–C). To evaluate whether pseudotime captured disease progression, we compared it with established neuropathological staging measures. Pseudotime was positively correlated with both Braak neurofibrillary tangle stage (Spearman ρ = 0.47, *p* = 9.3 × 10⁻¹³; *n* = 201; Fig. 2G) and Thal amyloid phase (Spearman ρ = 0.34, *p* = 1.1 × 10⁻⁵; Fig. 2H), demonstrating agreement between the inferred molecular trajectory and neuropathological progression. To determine whether pseudotime primarily captured disease progression rather than demographic heterogeneity, we examined its association with age, sex, and race. Among donors younger than 90 years, pseudotime was not associated with age at death (*p* = 0.37; Fig. 2D). The same lack of association was observed in the complete cohort (*p* = 0.51; Supplementary Fig. 2). Pseudotime also did not differ significantly by sex (*p* = 0.646; Fig. 2E) or between White and Black or African American donors (*p* = 0.708; Fig. 2F). Together, these findings indicate that the inferred pseudotime primarily reflects Alzheimer’s disease progression rather than demographic heterogeneity within this cohort.

**Fig. 2.**
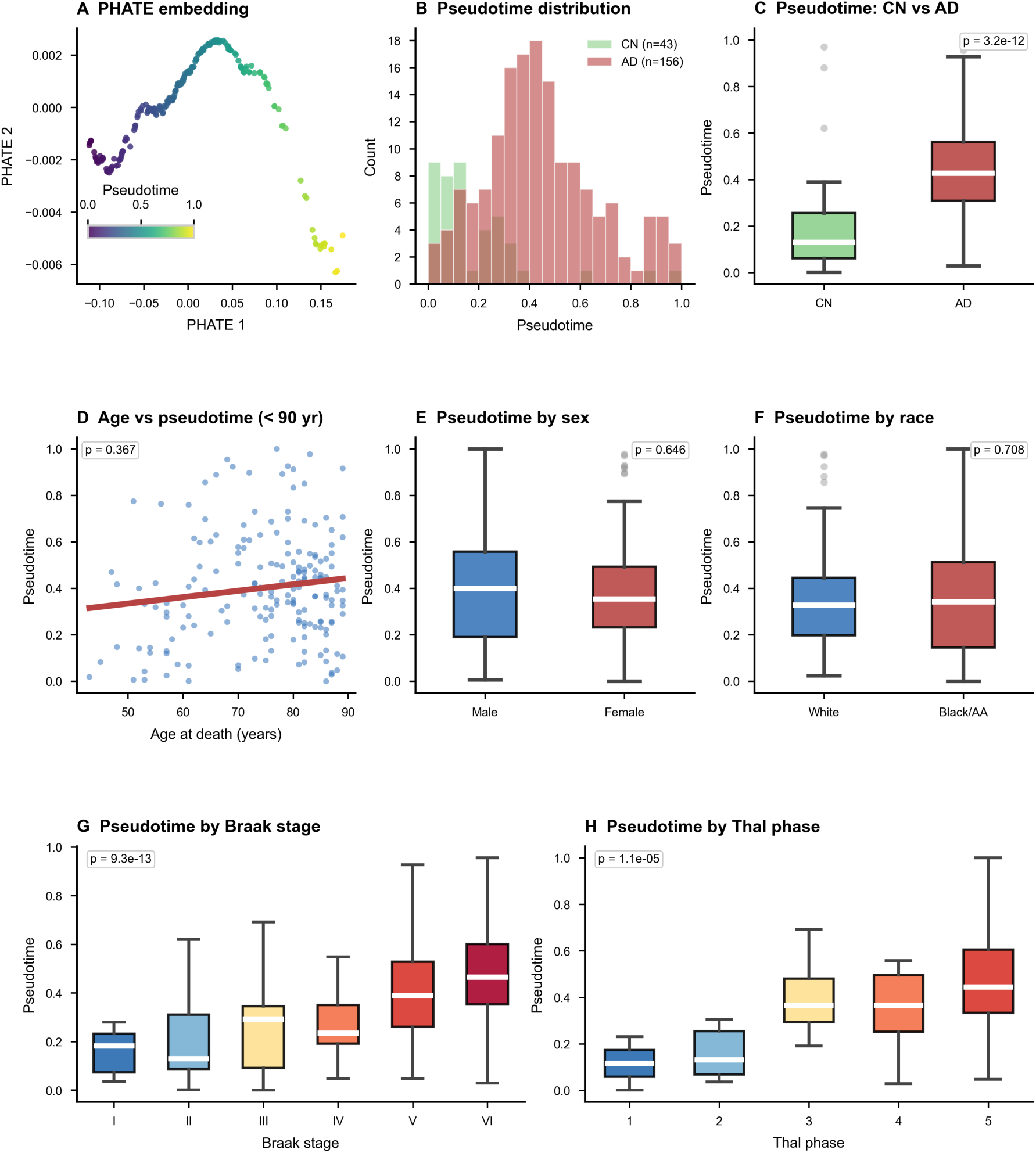
PHATE pseudotime recapitulates Alzheimer’s disease progression. (A) PHATE embedding of the cross-regional proteome (n = 213), colored by pseudotime. (B–C) Pseudotime was significantly higher in AD than in cognitively normal donors (Mann-Whitney U p = 3.2 × 10⁻¹²). (D) Pseudotime was not associated with age at death among donors <90 years (p = 0.367). (E–F) Pseudotime did not differ by sex (p = 0.646) or between White and Black or African American donors (p = 0.708). (G–H) Pseudotime correlated positively with Braak stage (Spearman ρ = 0.47, p = 9.3 × 10⁻¹³) and Thal phase (p = 1.1 × 10⁻⁵).

### Distinct trajectory clusters define regional proteomic progression programs

Trajectory clustering identified three protein clusters with distinct temporal trajectories and regional composition (Fig. 3A–C). All three clusters exhibited a major transition near pseudotime ≈0.6, which we refer to as the molecular tipping point. Cluster 0 (C0) comprised 2,377 high-confidence proteins and was almost exclusively represented by DLPFC proteins (2,312 DLPFC; 65 STG). These proteins remained relatively stable before declining after the tipping point (Fig. 3A). Cluster 1 (C1) contained 1,724 proteins and was similarly dominated by STG proteins (1,624 STG; 100 DLPFC). In contrast to C0, C1 proteins declined progressively before the tipping point and showed partial recovery at later pseudotime values (Fig. 3B). Cluster 2 (C2) comprised 3,211 proteins from both cortical regions, although STG proteins were more abundant (1,988 STG; 1,223 DLPFC). These proteins increased progressively before plateauing near the tipping point (Fig. 3C). The relatively small numbers of STG proteins assigned to C0 and DLPFC proteins assigned to C1 suggest that these minority assignments likely reflect weak cluster membership rather than distinct cross-regional trajectory programs. Protein-level cluster assignments, membership scores, and trajectory statistics are provided in Supplementary Table 2.

**Fig. 3.**
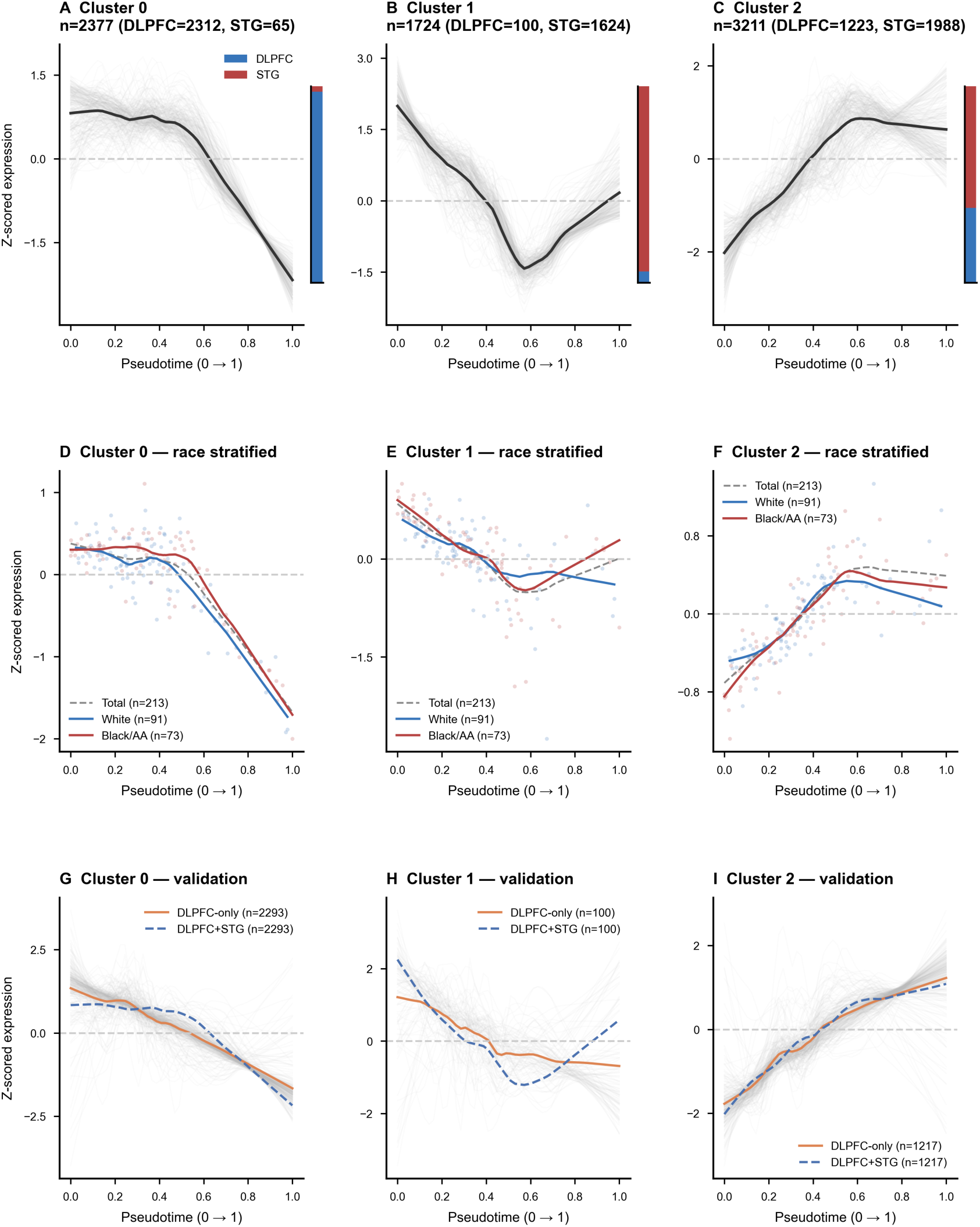
Three protein trajectory clusters with distinct regional compositions. (A–C) Fuzzy c-means clustering identified three high-confidence trajectory clusters (membership ≥0.99): DLPFC-enriched Cluster 0, STG-enriched Cluster 1 and Cluster 2 exhibiting coordinated cross-regional increase. (D–F) Race-stratified trajectories for 3 clusters were qualitatively consistent between White (n = 91) and Black or African American (n = 73) donors. (G–I) Trajectories derived from the non-overlapping DLPFC-only cohort are shown alongside the corresponding trajectories from the primary cross-regional analysis.

We next evaluated the robustness of these trajectory patterns across racial groups and in an independent validation cohort. The overall trajectory shapes of all three clusters were qualitatively similar between White and Black or African American donors (Fig. 3D–F), indicating that the major temporal patterns were preserved across these groups. Reproducibility was further evaluated in the independent DLPFC-only cohort. The average DLPFC trajectories of all three clusters closely resembled those observed in the matched cross-regional cohort (Fig. 3G–I). Reproducibility was strongest for C0 and C2, whereas C1 showed greater variability, consistent with the relatively small number of DLPFC proteins assigned to this predominantly STG-derived cluster.

### A subset of early declining STG proteins exhibits later decline in the DLPFC

All three trajectory clusters exhibited a major transition near pseudotime ≈0.6 (Fig. 3A–C), which we designated as the molecular tipping point and used to define early (PT 0–0.6) and late (PT 0.6–1.0) trajectory windows. Within the early trajectory window, STG Cluster 1 (C1) proteins declined progressively, whereas STG Cluster 2 (C2) proteins increased over the same interval (Fig. 4A). At the individual level, STG C1 and C2 principal component (PC1) scores were strongly anti-correlated (Spearman ρ = −0.77, P = 1.4 × 10⁻⁴²; Fig. 4B), indicating coordinated suppression of one molecular response accompanied by activation of another during early molecular progression. Protein-level regression analysis identified proteins exhibiting significant pseudotime-associated changes during early STG progression (Fig. 4C). The most strongly declining proteins included *CAP2, HPCA, PRKCE, ACTN2*, and *DLG3*, whereas *HSPB1*, *DBI*, and *DEK* were among the proteins showing the strongest increases. Among STG proteins in C1, 1,562 showed significant pseudotime-associated decline during PT 0–0.6 (FDR < 0.05). Among these, 169 proteins overlapped with DLPFC C0 proteins that exhibited significant decline during the later PT 0.6–1.0 interval (FDR < 0.05), indicating distinct regional timing (Fig. 4D). These proteins showed progressive decline in the STG during the early trajectory window but only minimal change in the DLPFC before PT 0.6, after which they declined progressively. Because none of the C1 proteins remained significantly associated with pseudotime beyond PT 0.6 (all FDR > 0.05), the STG trajectories are shown only within the early trajectory window. These shared proteins therefore represent a subset of the early STG decline program that is also observed in the DLPFC but during a later stage of molecular progression.

**Fig. 4.**
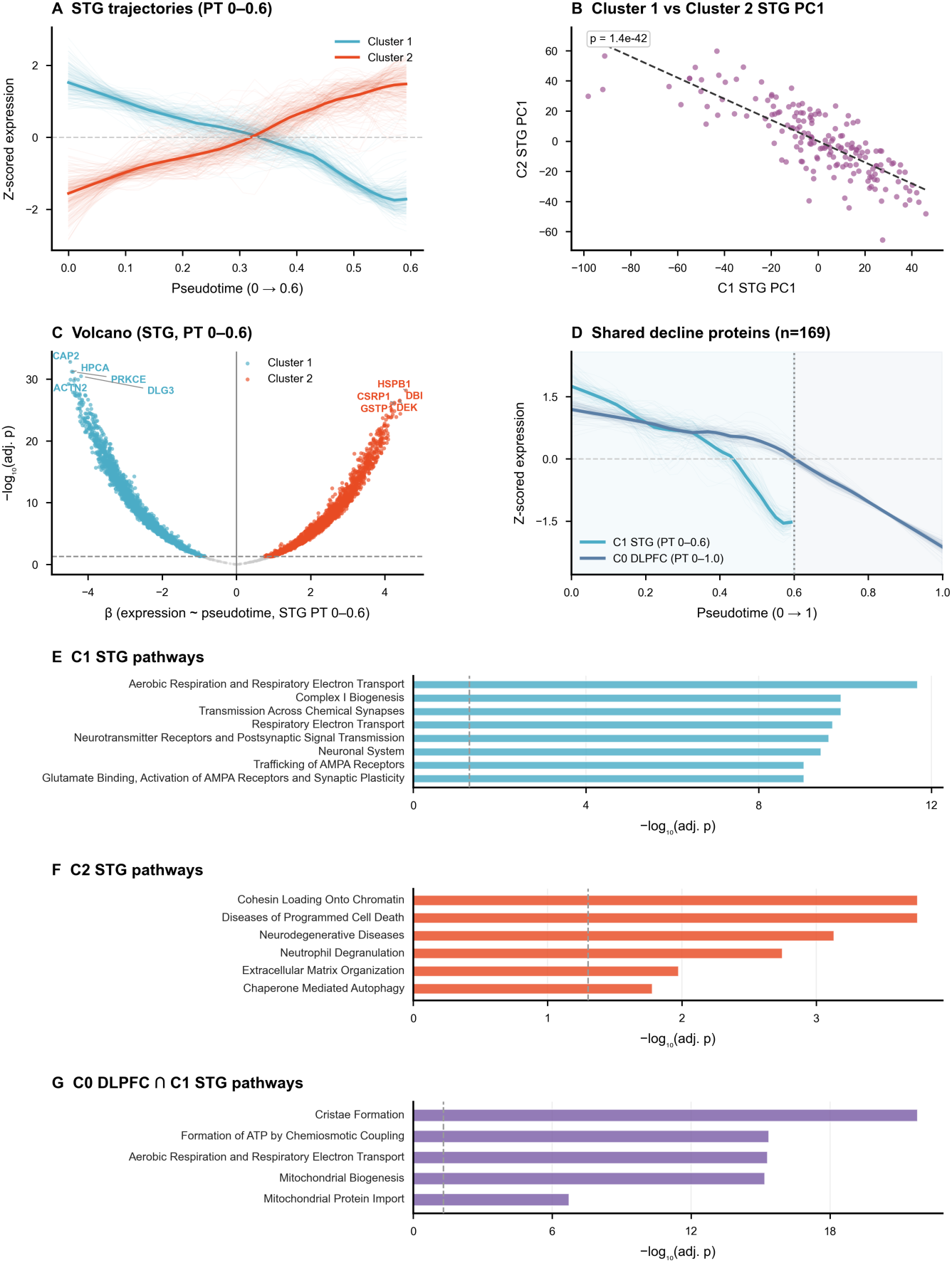
Proteomic dynamics within STG before the molecular tipping point. (A) STG Cluster 1 (C1) and Cluster 2 (C2) trajectories over pseudotime (PT) 0–0.6, showing C1 suppression and C2 activation. (B) C1 and C2 STG PC1 scores were inversely correlated in pre-tipping individuals (n = 174; Spearman’s ρ = −0.77, P = 1.4 × 10⁻⁴²). (C) Volcano plot of STG protein expression regressed on pseudotime, with selected significant proteins labeled. (D) Mean trajectories of 169 proteins shared between early decreasing C1 STG proteins during and decreasing C0 DLPFC proteins. The STG trajectory is shown only through PT 0.6 because none of the C1 proteins exhibited significant pseudotime association beyond this point (all FDR > 0.05). (E,F) Representative non-redundant Reactome pathways enriched among decreasing C1 and increasing C2 STG proteins, respectively. (G) Representative non-redundant Reactome pathways enriched among the 169 shared proteins. STG, superior temporal gyrus; DLPFC, dorsolateral prefrontal cortex.

Functional enrichment analysis indicated that C1 proteins were primarily associated with synaptic transmission and mitochondrial bioenergetic processes, whereas C2 proteins were enriched for pathways related to chromatin organization, programmed cell death, innate immune responses, extracellular matrix organization, and stress-associated signaling (Fig. 4E,F; Supplementary Table 3). Notably, the 169 proteins exhibiting distinct regional timing were specifically enriched for mitochondrial organization and energy metabolism (Fig. 4G, Supplementary Table 3), indicating that delayed DLPFC progression predominantly involved proteins associated with mitochondrial function. Together, these findings indicate that early STG progression is characterized by coordinated suppression of synaptic and mitochondrial protein programs together with activation of cellular pathways related to chromatin remodeling, stress signaling, and innate immune responses.

### A coordinated cross-regional protein program exhibits more rapid progression in the superior temporal gyrus

Cluster 2 included proteins from both the DLPFC (n = 1,223) and STG (n = 1,988) that increased progressively before the molecular tipping point (pseudotime 0–0.6). The average trajectories of C2 proteins closely tracked each other across the two regions (Fig. 5A). This demonstrates coordinated temporal dynamics of the C2 program across both cortical regions. Consistent with this observation, C2 principal component (PC1) scores were significantly correlated between the DLPFC and STG across pre-tipping individuals (Spearman ρ = 0.52, *p* = 5.5 × 10⁻¹³; Fig. 5B). Individuals with stronger activation of the C2 program in one region also exhibited stronger activation in the other.

**Fig. 5.**
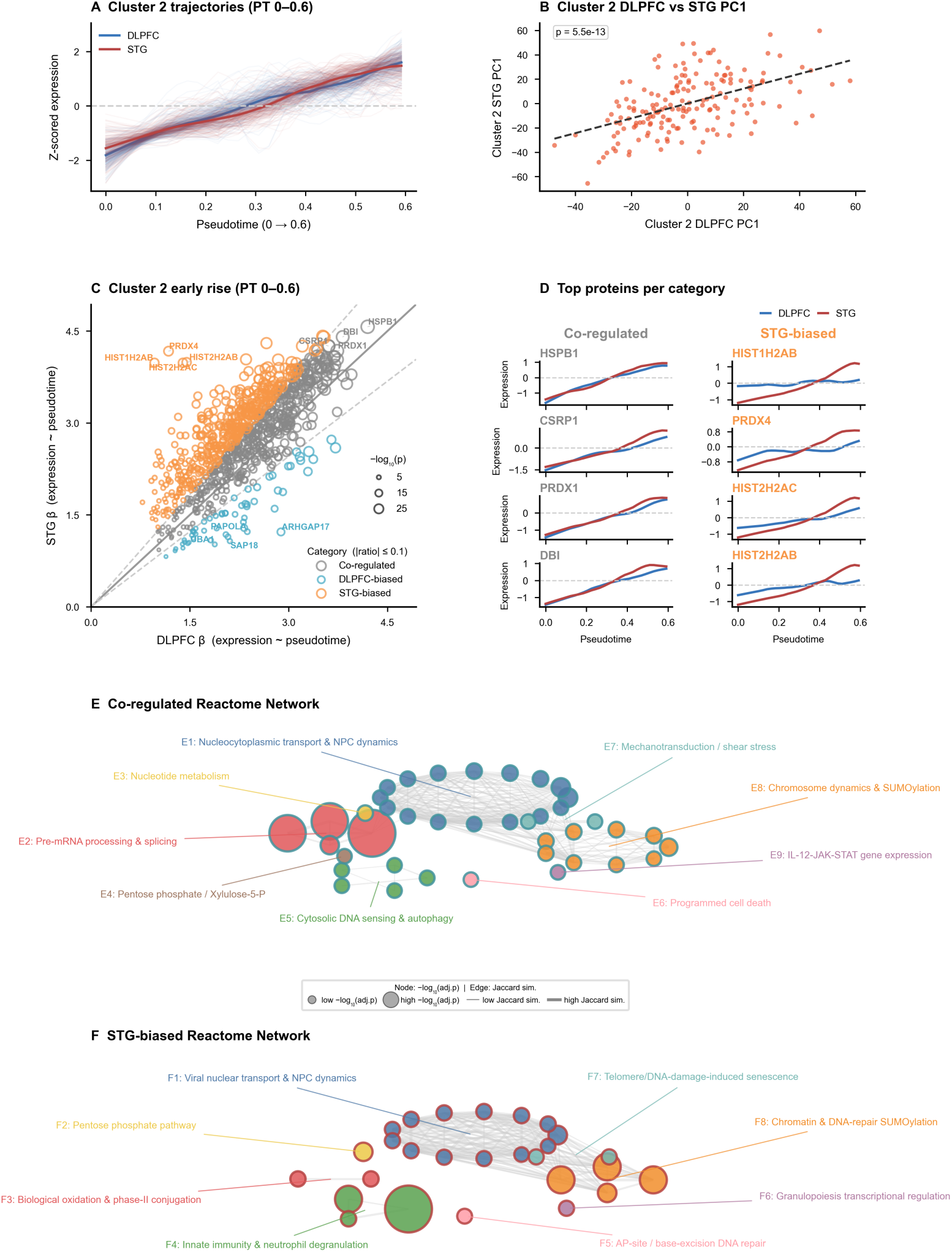
Coordinated DLPFC–STG dynamics within Cluster 2. (A) Cluster 2 protein trajectories in DLPFC (n = 1,223) and STG (n = 1,988) over pseudotime < 0.6. (B) Cluster 2 DLPFC and STG PC1 scores were positively correlated among pre-tipping individuals (Spearman ρ = 0.52, p = 5.5 × 10⁻¹³). (C) DLPFC versus STG regression coefficients (β) for Cluster 2 proteins, categorized as co-regulated, DLPFC-biased, or STG-biased using asymmetry index. (D) Trajectories for representative co-regulated and STG-biased proteins. (E–F) Reactome pathway networks for co-regulated and STG-biased Cluster 2 proteins.

Although C2 proteins shared similar trajectory shapes, they did not necessarily exhibit the same rate of increase along pseudotime. Trajectory clustering was performed on row-wise standardized trajectories, grouping proteins according to their temporal pattern rather than the magnitude of change. We therefore compared protein-specific regression coefficients between the DLPFC and STG to quantify regional differences in the rate of change. Many proteins clustered close to the diagonal (Fig. 5C), indicating broadly comparable temporal dynamics between the two regions. However, the overall distribution was shifted above the diagonal, with substantially more proteins exhibiting larger regression coefficients in the STG than in the DLPFC. Thus, while C2 proteins were largely co-upregulated across both regions, their rates of increase were generally greater in the STG. Representative examples illustrate these two components of the C2 program. HSPB1, CSRP1, PRDX1, and DBI displayed nearly identical trajectories in the DLPFC and STG. In contrast, HIST1H2AB, HIST2H2AB, HIST2H2AC, and PRDX4 exhibited substantially steeper increases in the STG (Fig. 5D). Only a small number of proteins showed modestly larger regression coefficients in the DLPFC.

Functional enrichment analysis further distinguished these two protein groups. Co-regulated proteins were enriched for RNA metabolism, including mRNA splicing and transcript transport, together with SUMOylation, cytokine/interferon signaling, and cell-cycle-related processes (Fig. 5E). In contrast, proteins with greater rates of increase in the STG were preferentially enriched for innate immune pathways (including neutrophil degranulation), biological oxidations, SUMOylation-associated DNA damage responses, and nuclear transport (Fig. 5F). Complete pathway enrichment results for the co-regulated, STG-biased, and DLPFC-biased protein sets are provided in Supplementary Table 3.

### Superior temporal gyrus precedes DLPFC along the shared cross-regional C2 protein program

Although C2 proteins increased concordantly across the DLPFC and STG, the two regions did not necessarily follow this rising trajectory at the same pseudotime position for each protein. We therefore examined the relative timing at which individual C2 proteins crossed a common trajectory landmark in the two brain regions. For each C2 protein, crossing time was defined as the first pseudotime point at which the smoothed, Z-scored trajectory exceeded zero for at least three consecutive grid points, corresponding to a transition from below-average to above-average relative expression on that protein’s normalized trajectory. Among 931 C2 proteins with measurable crossing times in both the DLPFC and STG, 74% (689/931) crossed earlier in the STG than in the DLPFC (ΔPT = STG crossing time − DLPFC crossing time; one-sided Wilcoxon signed-rank test, p = 7.95 × 10⁻⁷⁵; Fig. 6A,B), indicating a systematic temporal lead of the STG. This regional lead was not uniform across C2 protein groups. STG-biased proteins showed the largest negative ΔPT values, whereas co-regulated proteins displayed smaller regional offsets and DLPFC-biased proteins showed a more heterogeneous pattern (Fig. 6C). Although DLPFC-biased proteins exhibited larger regression coefficients in the DLPFC, they also showed a modest STG lead, indicating that regional differences in the rate of progression and the timing of progression capture complementary aspects of trajectory dynamics. Thus, proteins with stronger STG pseudotime effects also tended to reach the same trajectory landmark earlier in the STG.

**Fig. 6.**
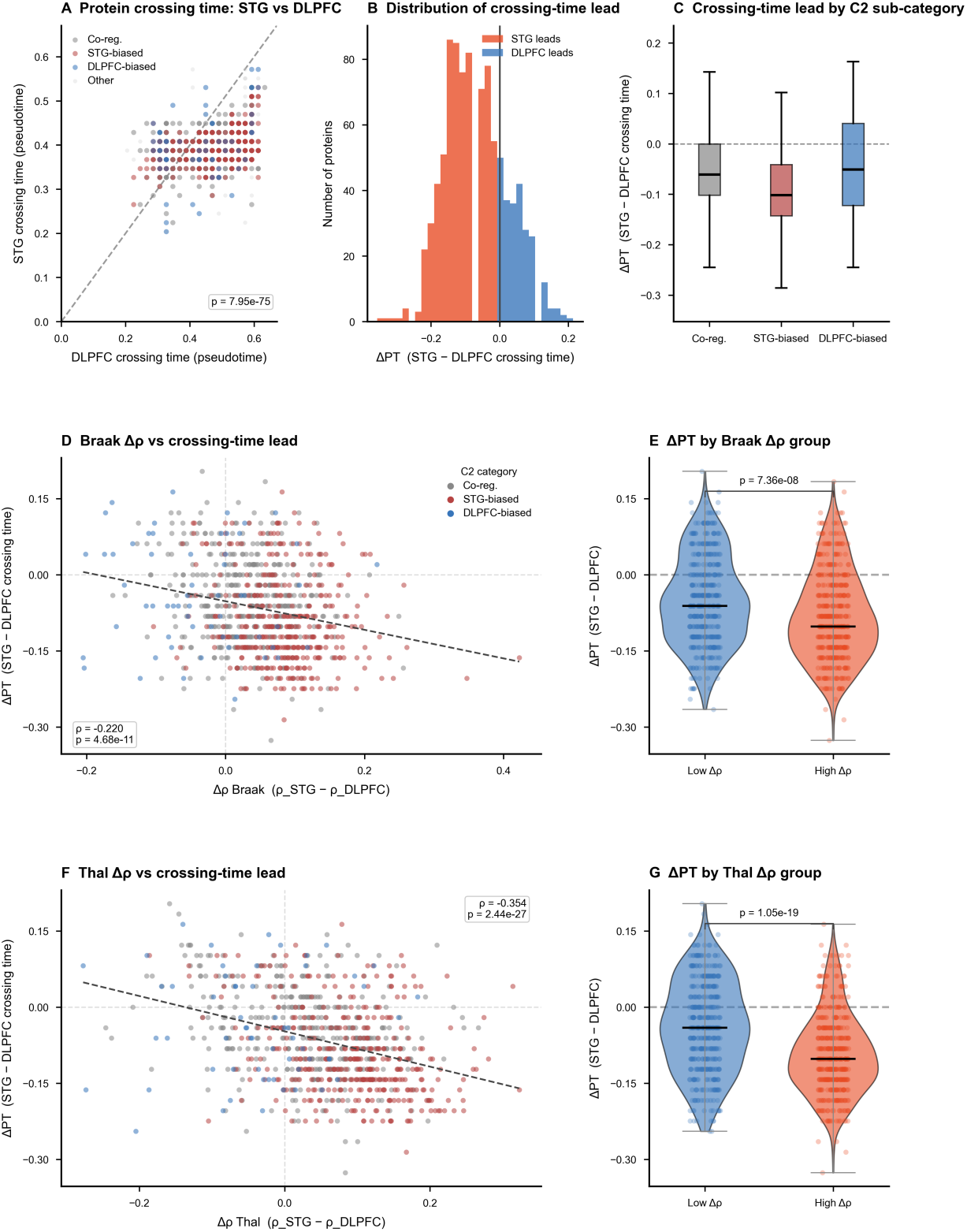
The superior temporal gyrus precedes the DLPFC along the shared Cluster 2 trajectory. (A) STG versus DLPFC activation (crossing) time for Cluster 2 proteins. STG activation was significantly earlier than DLPFC activation (one-sided Wilcoxon signed-rank p = 7.95 × 10⁻⁷⁵), with 74% of proteins showing an STG lead. (B) Distribution of activation-time lead (ΔPT = STG − DLPFC crossing time): STG-leading (n = 689) versus DLPFC-leading (n = 242) proteins. (C) ΔPT by Cluster 2 sub-category (co-regulated, STG-biased, DLPFC-biased). The activation-time lead was most substantial among STG-biased proteins. (D) *Δρ_Braak_* (regional difference in protein–Braak stage correlation) versus ΔPT showed strong negative correlation (Spearman ρ = −0.220, p = 4.68 × 10⁻¹¹). (E) ΔPT distribution for low-versus high-*Δρ_Braak_* proteins (median split); high-*Δρ_Braak_* proteins showed a stronger STG lead in activation time (Mann-Whitney U p = 7.36 × 10⁻⁸). This indicates that proteins with stronger Braak-association in STG will likely activate earlier in STG. (F) *Δρ_T_*_ℎ*al*_ (regional difference in protein–Thal phase correlation) versus ΔPT; negatively correlated (Spearman ρ = −0.354, p = 2.44 × 10⁻²⁷). (G) ΔPT for low-versus high-*Δρ_T_*_ℎ*al*_ proteins; high-*Δρ_T_*_ℎ*al*_ proteins showed a stronger STG lead (p = 1.05 × 10⁻¹⁹).

To determine whether this STG temporal lead was related to neuropathologic progression, we compared the regional association of each C2 protein with Braak stage and Thal amyloid phase. For each protein, we computed Δρ_Thal = ρ_STG − ρ_DLPFC, where ρ_STG and ρ_DLPFC represent the Spearman correlations between regional protein expression and donor-level Thal phase in the STG and DLPFC, respectively. Δρ_Braak was defined analogously using Braak neurofibrillary tangle stage. A positive Δρ indicates that the protein’s expression is more strongly associated with donor-level neuropathologic stage in the STG than in the DLPFC. Across C2 proteins, Δρ_Thal was significantly negatively correlated with ΔPT (Spearman ρ = −0.354, p = 2.44 × 10⁻²⁷; Fig. 6F), indicating that proteins with stronger STG-than-DLPFC association with Thal phase tended to cross earlier in the STG. A similar relationship was observed for Braak stage (Δρ_Braak; Spearman ρ = −0.220, p = 4.68 × 10⁻¹¹; Fig. 6D).

To further evaluate the relationship between regional progression and neuropathology, proteins were stratified into high and low Δρ groups based on the median regional difference in pathology association. Proteins showing stronger association with Thal phase in the STG than in the DLPFC (larger Δρ_Thal values) exhibited significantly more negative ΔPT values than proteins in the low Δρ_Thal group (Mann–Whitney U *P* = 1.05 × 10⁻¹⁹; Fig. 6G). A similar relationship was observed for Braak stage (Mann–Whitney U *P* = 7.36 × 10⁻⁸; Fig. 6E). These findings indicate that proteins reaching the shared trajectory earlier in the STG also tend to exhibit stronger associations with amyloid and tau neuropathology in the STG than in the DLPFC. Complete protein-level crossing times, regional pathology associations, and differential pathology statistics are provided in Supplementary Table 4.

## Discussion

Previous multi-region proteomic studies established that Alzheimer’s disease (AD) comprises both shared and region-specific molecular programs across vulnerable brain regions^5,6,8^. Our study builds upon these observations by incorporating the temporal dimension of molecular progression. By reconstructing continuous proteomic trajectories from matched dorsolateral prefrontal cortex (DLPFC) and superior temporal gyrus (STG), we identified two complementary patterns of cross-regional progression. Proteins involved in mitochondrial bioenergetics declined substantially earlier in the STG than in the DLPFC, whereas a second protein program increased in both regions throughout disease progression but generally progressed more rapidly in the STG. Notably, the inferred progression axis was independent of age, sex, and race in this diverse cohort, supporting the interpretation that it primarily captures molecular disease progression rather than demographic heterogeneity. Together, these findings demonstrate that molecular programs shared across vulnerable cortical regions can exhibit distinct regional temporal progression during AD.

Within the STG, suppression of synaptic and mitochondrial protein programs was accompanied by coordinated activation of pathways related to chromatin remodeling, cellular stress signaling, innate immune responses, and extracellular matrix organization. This coordinated organization is consistent with previous network-based proteomic analyses demonstrating coordinated reductions in neuronal and synaptic modules together with increases in glial- and immune-associated molecular programs in the AD brain^5,7^. The agreement between our trajectory analysis and previous co-expression studies indicates that the inferred progression faithfully recapitulates established molecular organization within the temporal cortex. Across cortical regions, however, a different pattern emerged. A subset of proteins involved in mitochondrial bioenergetics followed temporally offset trajectories across cortical regions, declining substantially later in the DLPFC than in the STG. These proteins were enriched for mitochondrial bioenergetics and oxidative phosphorylation, indicating that disruption of energy metabolism progresses with markedly different timing across vulnerable cortical regions. Large-scale proteomic studies have consistently identified disruption of mitochondrial and energy metabolism as a core feature of the AD brain. Johnson et al. identified early alterations in proteins involved in energy metabolism together with glial activation in brain tissue and cerebrospinal fluid^4^, while subsequent deep proteomic analyses further demonstrated widespread dysregulation of mitochondrial and metabolic pathways across the AD brain^5^. Our findings further indicate that proteins involved in mitochondrial bioenergetics are shared across vulnerable cortical regions but unfold with distinct regional timing. This regional sequence is consistent with FDG-PET studies demonstrating that cerebral glucose hypometabolism develops earlier in temporal cortical regions than in frontal cortical regions during AD progression^9,10^. Although our cross-sectional analysis cannot directly establish sequential regional progression, the similarity between proteomic trajectories and metabolic imaging suggests that temporally offset mitochondrial dysfunction may contribute to the well-established temporal-to-frontal sequence of metabolic impairment in AD.

The second major finding was the identification of a broadly coordinated protein program that increased across both cortical regions. Most proteins within this trajectory exhibited similar temporal profiles in the DLPFC and STG and were enriched for RNA metabolism, mRNA processing, SUMOylation, cytokine signaling, and protein homeostasis. These biological processes are increasingly recognized as fundamental components of the cellular response to proteotoxic stress and neurodegeneration^11–13^. Previous cross-regional proteomic studies demonstrated that many of these molecular responses are shared across multiple vulnerable brain regions in AD^6,8^. Despite this overall coordination, regional differences remained evident. Across most proteins, increases along the shared trajectory were generally more rapid in the STG than in the DLPFC, although these regional differences were modest for most proteins. The largest regional differences were observed for only a small subset of proteins. These proteins exhibited more rapid increases along the shared trajectory, reached comparable proteomic states earlier, and showed stronger associations with donor-level Braak stage and Thal phase in the STG. Together, these findings demonstrate that molecular responses shared across vulnerable cortical regions are not necessarily synchronized but instead can differ in their temporal progression. Rather than identifying region-specific molecular programs, our trajectory analysis suggests that shared molecular responses can reach comparable proteomic states at different stages of disease progression across cortical regions. This interpretation is consistent with the earlier neuropathological involvement of temporal cortical regions relative to frontal regions in AD^14^.

Several limitations should be considered. First, the inferred molecular progression was reconstructed from cross-sectional rather than longitudinal data and therefore represents an approximation of disease progression at the population level, a limitation inherent to current postmortem human brain studies. Second, like most postmortem AD cohorts, the AMP-AD Diverse Cohorts dataset is enriched for donors with advanced Braak stages. Consequently, the inferred trajectories provide limited resolution of the earliest molecular changes during AD progression. Third, although the major DLPFC trajectory patterns were reproduced in an independent cohort, validation of the cross-regional temporal organization in an independent dataset with matched DLPFC and STG proteomes will be important to establish the robustness of these findings. Future studies using independent matched multi-region cohorts will determine whether the temporal relationships identified here are conserved across additional AD-vulnerable brain regions.

In summary, our study extends previous cross-regional proteomic analyses by incorporating the temporal dimension of molecular progression. Rather than identifying additional disease-associated pathways, our findings demonstrate that established molecular programs exhibit distinct regional temporal organization across vulnerable cortical regions. These findings suggest that regional vulnerability in AD is reflected not only in the molecular programs involved but also in their temporal organization during disease progression.

## Methods

### Study cohort and data

The AMP-AD Diverse Cohorts study is a cross-consortium initiative generating harmonized postmortem brain genomic, transcriptomic, and proteomic data from a racially and ethnically diverse population. Brain tissue was contributed by multiple institutions, including Mayo Clinic, Rush University, Mount Sinai University Hospital, and Emory University, and was processed centrally for proteomic profiling^6^. Neuropathological data across contributing sites were harmonized to common diagnostic criteria, including Braak staging and CERAD scoring, prior to data release^6^. TMT proteomics data from this study were accessed from the AD Knowledge Portal (Synapse)^15^. Data were available for the DLPFC (n = 1,086 specimens from 579 individual donors) and STG (n = 278 specimens from 242 individual donors). Donors with proteomic data from both regions were identified for the primary cross-regional cohort; donors with phenotype data missing required annotations, or failing quality-control criteria, were excluded from the analytic pseudotime cohort. A non-overlapping DLPFC-only cohort was used for validation.

### Data preprocessing and quality control

TMT-MS proteomic quantification, batch-effect correction, and initial quality control, including removal of proteins with more than 50% missing values and iterative principal component analysis to exclude outliers within each region, were performed as previously described for this cohort^6^. Individuals with more than 50% missing protein values were excluded. Remaining missing values were imputed by column-wise median. Expression values were log-transformed and Z-score normalized per sample. For cross-regional analysis, DLPFC and STG protein vectors were concatenated per donor, yielding 18,036 region-annotated features across 9,018 unique proteins. Thus, the same protein measured in different brain regions was treated as two region-specific features, enabling direct comparison of their proteomic trajectories across regions. Covariate effects of age at death, sex, and postmortem interval (PMI) were removed by ordinary least squares regression. Residuals were retained for trajectory analysis.

### PHATE embedding and pseudotime assignment

Principal component analysis (PCA) was applied to the residualized, cross-regional expression matrix. Components were retained up to the number explaining 90% or more of the total variance, with the number of retained components capped at 100. The reduced matrix was embedded in two dimensions using PHATE (Potential of Heat-diffusion for Affinity-based Trajectory Embedding)^16^, with k-nearest neighbors set to 10, diffusion time t set to 100, and a fixed random seed of 42. The resulting embedding formed a smooth trajectory curve along which donors were ordered. For each donor, pseudotime was calculated as the normalized geodesic distance along this curve from its control-enriched end, scaled to a range of 0 to 1. Donors with similar cross-regional proteomic profiles were expected to lie near one another on this curve and therefore share similar pseudotime values. This pseudotime was accordingly treated as a continuous quantification of proteomic dynamics along Alzheimer’s disease progression. Derived pseudotime was validated against Braak neurofibrillary tangle stage and Thal amyloid phase using Spearman correlation, and against AD diagnosis using a Mann-Whitney U test.

### Trajectory-based clustering of protein dynamics

For each protein in each region, expression was smoothed against pseudotime using locally weighted scatterplot smoothing (LOWESS, span = 0.30)^17^ evaluated at 50 evenly spaced pseudotime grid points. Trajectories were row-wise Z-scored prior to clustering to ensure comparability across proteins with different expression scales.

Smoothed, Z-scored trajectories from both regions were jointly clustered using fuzzy c-means with soft dynamic time warping (soft-DTW, *γ* = 0.01)^18^ as the distance metric, implemented via the tslearn Python library^19^. We experimented with different number of clusters K = 2–10, and selected K = 3 based on the point of inflection with the minimum inter-centroid distance curve (Supplementary Fig. 3). Membership scores were computed via a fuzzy distance-to-membership transformation (fuzziness m = 2.0). Distances to cluster centroids were converted to fuzzy memberships using the standard fuzzy c-means update rule with m = 2.0. High-confidence cluster assignments require a stringent membership score of 0.99 or greater.

### Race-stratified trajectory analysis

For each cluster, LOWESS-smoothed protein trajectories were computed separately within White and Black or African American donors in the primary cross-regional cohort, using the same smoothing parameters as the primary analysis. Donors classified as other race/ethnicity were not analyzed separately due to their smaller sample size. Race-stratified trajectories were compared with the corresponding trajectory computed across the full analytic cohort to assess visual consistency in progression trajectory shape.

### DLPFC-only cohort validation

The PHATE and trajectory-smoothing pipeline was applied independently to the non-overlapping DLPFC-only cohort (n = 364), using preprocessing, embedding, and smoothing parameters identical to those used in the primary cross-regional analysis. For each cluster, individual protein trajectories were derived and averaged in the same manner as in the primary analysis to obtain a representative DLPFC-only trajectory. These trajectories were then compared with the one derived from primary cross-regional cohort to assess reproducibility.

### Cluster-level co-expression

Across all 3 protein clusters we observed a consistent transition at approximately pseudotime 0.6 (Fig. 3A–C). This shared transition point was designated the tipping point and used to divide individuals into pre-tipping (pseudotime < 0.6; n = 174) and post-tipping (pseudotime ≥ 0.6; n = 39) groups. As donors in this cohort are highly skewed toward advanced Braak and Thal stages (Fig. 1G–H), the pre-tipping window encompasses the majority of donors and spans a substantial range of neuropathological severity, whereas the post-tipping window comprises a smaller subset of donors at the far end of the trajectory. Subsequent analyses of within-region and cross-regional coordination were accordingly restricted to the pre-tipping window, in which sample size was adequate to support individual-level comparisons.

For each cluster-region combination, the first principal component (PC1) of high-confidence protein expression was computed across pre-tipping individuals. Spearman correlations between PC1 scores of different cluster-region pairs were computed within the pre-tipping group to assess coordinated or divergent dynamics. Linear regression of expression on pseudotime (PT 0–0.6) was performed per cluster per region to identify significant proteins (FDR-adjusted p < 0.05, Benjamini-Hochberg)^20^.

### Cross-regional rate comparison

Cluster 2 (C2) is the only cluster with substantial protein representation in both the DLPFC and STG. For each C2 protein significant in both regions, we evaluated the association of each protein with pseudotime in each brain region, yielding two regression coefficients for each protein: β_STG for association in STG and β_DLPFC for association in DLPFC. An asymmetry index was defined as |β_STG − β_DLPFC| / (|β_STG| + |β_DLPFC|). Proteins with an asymmetry index of 0.10 or less were classified as co-regulated. Remaining proteins were classified as STG-biased or DLPFC-biased according to which region showed the larger regression coefficient.

### Pathway enrichment and network visualization

Enrichment analysis was performed with gseapy^21^ using Enrichr^22^ against Reactome_Pathways_2024^23^, GO_Biological_Process_2025^24^, and KEGG_2026^25^. The background gene set comprised all proteins detected in the QC-passed expression matrix. For each queried protein set, pathways with an adjusted p < 0.05 were considered significantly enriched. For visualization, representative non-redundant Reactome pathways were selected from the significantly enriched pathways to summarize the major biological themes of each protein set, whereas complete enrichment results are provided in Supplementary Table 3. The Jaccard index was used to assess gene-set overlap between pathways, and communities of pathways were identified from this overlap using the Louvain algorithm^26^.

### Cross-regional comparison of protein activation time

For each high-confidence C2 protein present in both the DLPFC and STG, the protein was considered activated at a pseudotime point where its expression transitioned from below-average to above-average relative to its own trajectory. This crossing time (or activation time) was defined as the first pseudotime grid point at which the LOWESS-smoothed, Z-scored trajectory exceeded zero for at least three consecutive grid points. Proteins without a sustained crossing in a given region were excluded from that region’s crossing time. The temporal difference, ΔPT, was computed as STG crossing time minus DLPFC crossing time; a negative ΔPT indicates that the STG activates first. A Wilcoxon signed-rank test (H₁: median ΔPT < 0), with zeros excluded, was applied to all C2 proteins with valid crossing times in both regions (n = 931). Mann-Whitney U tests compared ΔPT distributions across the co-regulated, STG-biased, and DLPFC-biased sub-categories of proteins.

### Differential regional association with pathology stage

For each high-confidence protein in cluster 2, Spearman correlation was computed between its STG expression and Braak stage across all QC-passed individuals with available Braak staging, and separately between its DLPFC expression and Braak stage. The difference between these two correlations, denoted *Δρ_Braak_*, quantified how much more strongly (or weakly) the protein’s expression tracked Braak stage in the STG relative to the DLPFC. The same procedure was repeated using Thal phase in place of Braak stage, yielding *Δρ_T_*_ℎ*al*_. Finally, Spearman correlation between Δρ and ΔPT assessed the association between this regional differential and activation-time lead. Proteins were stratified by high versus low Δρ (median split), and Mann-Whitney U tests compared the resulting ΔPT distributions across these strata.

### Statistics

All analyses were performed in Python 3.10. Spearman correlations assessed ordinal associations; Mann-Whitney U tests compared between-group pseudotime distributions. Linear regression (OLS) estimated protein-pseudotime associations with FDR correction via the Benjamini-Hochberg procedure. Unless otherwise specified, tests were two-sided with a significance threshold of 0.05. The crossing-time lead test in Fig. 6A used a pre-specified one-sided Wilcoxon signed-rank test testing whether STG crossing times preceded DLPFC crossing times. Median-split *Δρ* group comparisons in Fig. 6E and Fig. 6G used the direction specified in the corresponding figure legends.

### Data and code availability

TMT proteomics data from the AMP-AD Diverse Cohorts study are available through the AD Knowledge Portal (https://adknowledgeportal.synapse.org) under Synapse ID syn53420674. Access to these data requires registration and approval of a Data Use Certificate in accordance with AMP-AD data governance policies. Braak stage, Thal phase, and other clinical and neuropathological metadata used in this study are available from the same resource.

## Acknowledgements

The results published here are in whole or in part based on data obtained from the AD Knowledge Portal Diverse Cohort Study (https://doi.org/10.7303/syn53420674). Data generation was supported by the following NIH grants: U01AG046139, U01AG046170, U01AG061357, U01AG061356, U01AG061359, and R01AG067025. We thank the donors and their families, and the investigators at Mayo Clinic, Rush University, Mount Sinai University Hospital, and Emory University, for their contributions to this dataset.

## Author contributions

T.G. conceived and designed the study, curated and analyzed the data, developed the methodology, interpreted the results, prepared the figures, and wrote the original manuscript. Y.U. and Y.P. assisted with pathway visualization. K.N. and A.S. applied for the data. J.Y. supervised the study, provided guidance on study design and interpretation, and reviewed and edited the manuscript. All authors read and approved the final manuscript.

