## Supplemental figure 1-3, Supplemental Table 1 for "Cross-Regional Trajectory Analysis Reveals Distinct Temporal Progression of Shared Proteomic Programs in Alzheimer’s Disease"

### Supplementary Information — Contents

**Supplementary Figure Legends** pp. 3–5

Supplementary Fig. 1 p. 3
PHATE embedding by demographic, diagnostic, and neuropathologic variables

Supplementary Fig. 2 p. 4
Age at death versus pseudotime in the full analytic cohort

Supplementary Fig. 3 p. 5
Cluster number selection for fuzzy c-means clustering

**Supplementary Table 1** p. 6
Cohort characteristics of the primary and validation arms

**Supplementary Table 2** —
Protein quantification and quality metrics (Excel workbook)

**Supplementary Table 3** —
Pathway enrichment results (Excel workbook)

**Supplementary Table 4** —
C2 protein crossing times and pathology associations (Excel workbook)

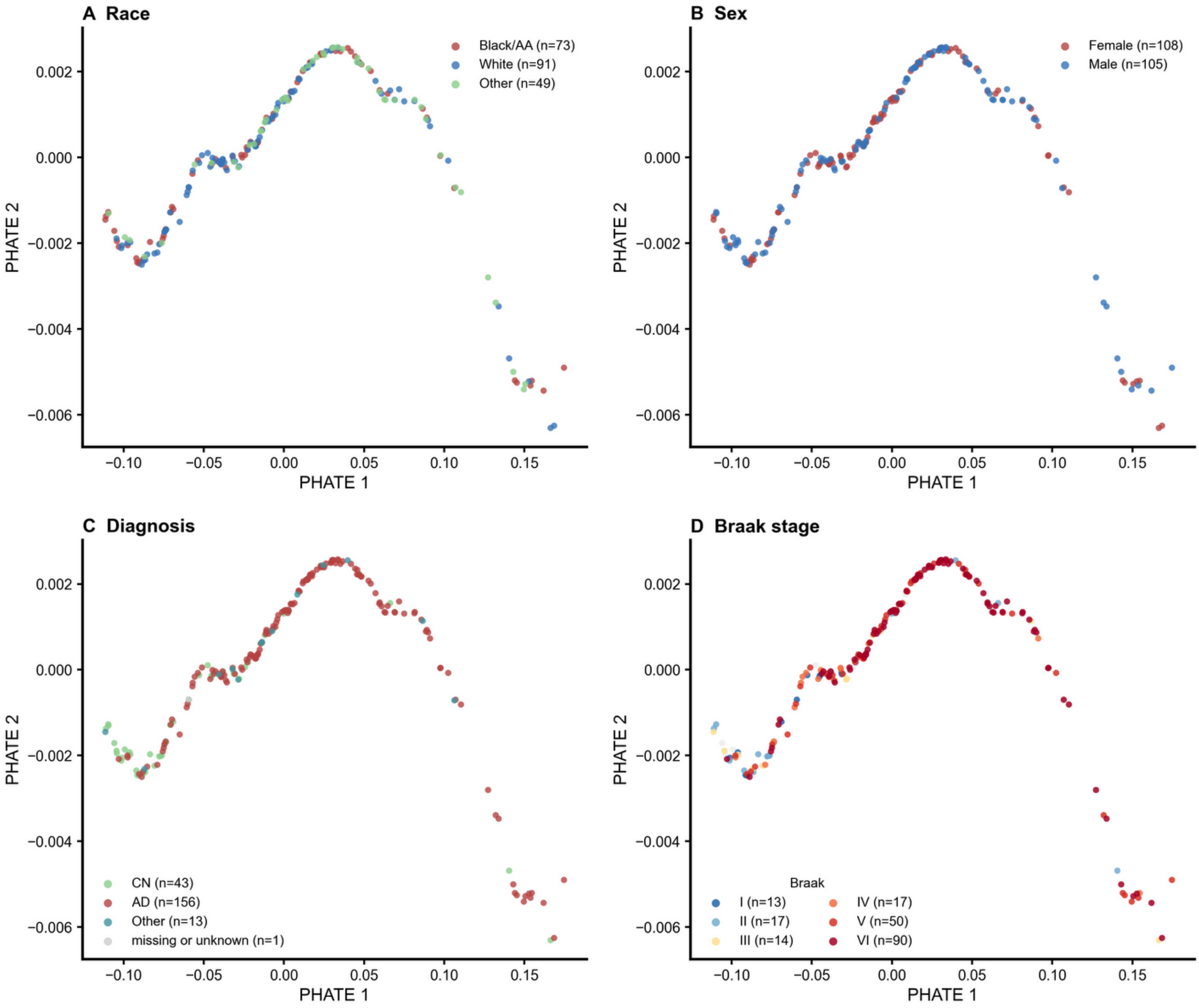


**Supplementary Fig. 1 |**

PHATE embedding colored by demographic, diagnostic, and neuropathologic variables. No systematic separation by any variable was observed along the PHATE manifold.

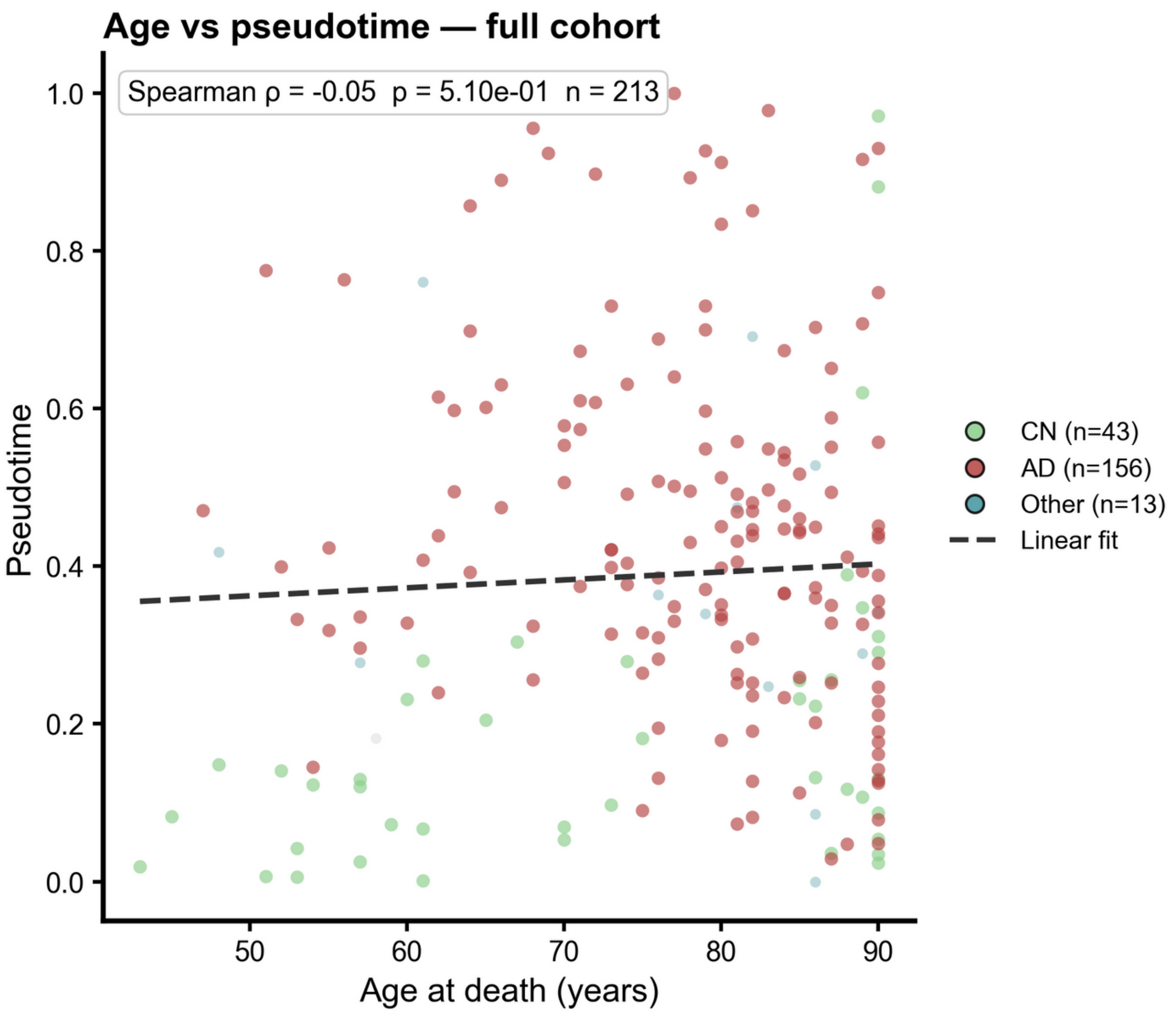


**Supplementary Fig. 2 |**

Age at death versus pseudotime in the full analytic cohort. Scatter plot of age at death versus pseudotime, colored by diagnosis, with ordinary least-squares fit. No significant association was observed.

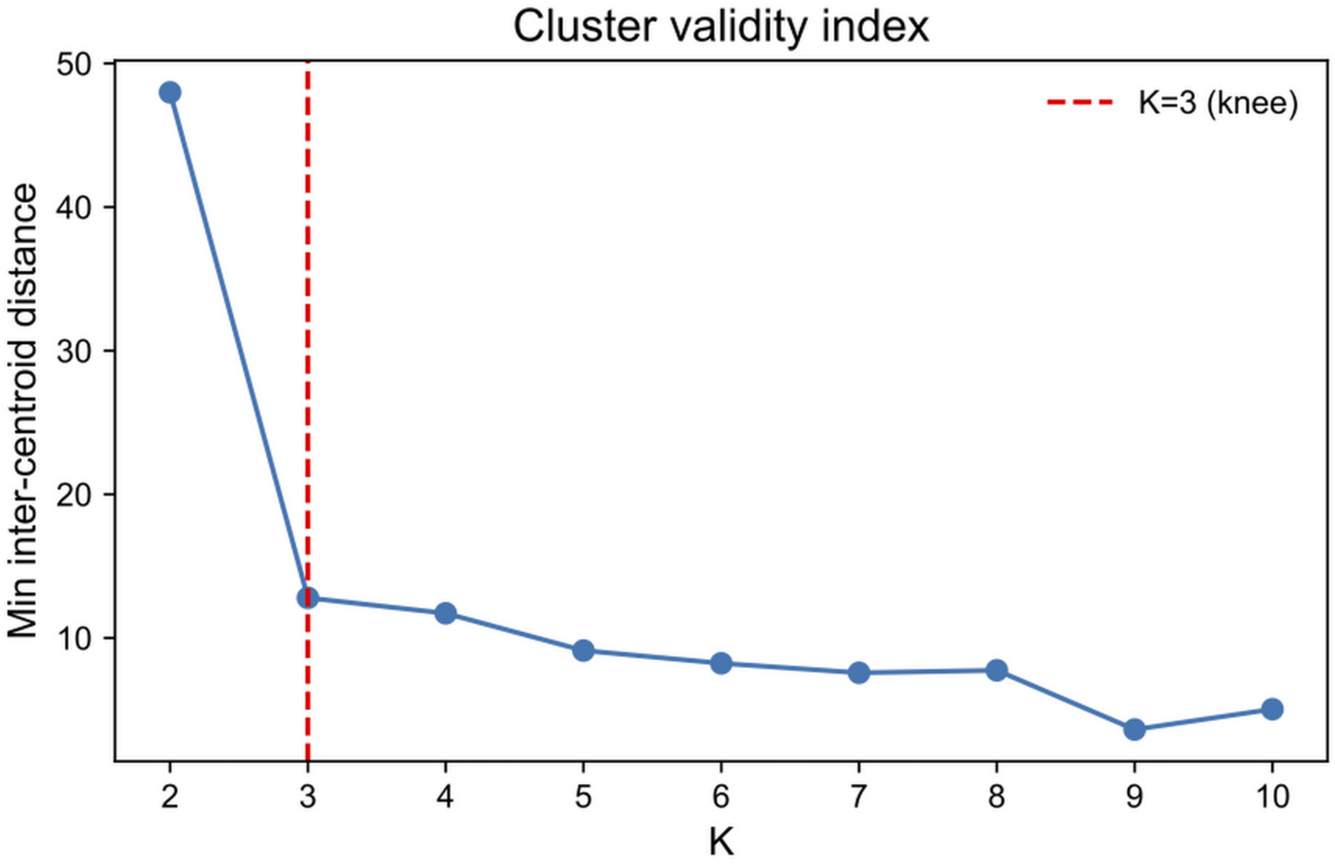


**Supplementary Fig. 3 |**

Cluster number selection. The minimum inter-centroid distance is plotted across candidate cluster numbers for fuzzy c-means clustering. K = 3 was selected based on the point of inflection in this curve.

**Supplementary Table 1. Cohort characteristics of the primary and validation arms**

| Variable | Sub-category | Primary arm (DLPFC+STG, n=213) | Validation arm (DLPFC-only, n=359) | Total (n=572) |
|---|---|---|---|---|
| n | | 213 | 359 | 572 |
| Diagnosis | AD | 156 (73.2%) | 191 (53.2%) | 347 (60.7%) |
| Diagnosis | Control | 43 (20.2%) | 64 (17.8%) | 107 (18.7%) |
| Diagnosis | Other | 13 (6.1%) | 103 (28.7%) | 116 (20.3%) |
| Diagnosis | Unknown | 1 (0.5%) | 1 (0.3%) | 2 (0.3%) |
| Sex | Male | 105 (49.3%) | 146 (40.7%) | 251 (43.9%) |
| Sex | Female | 108 (50.7%) | 213 (59.3%) | 321 (56.1%) |
| Race/Ethnicity | White | 91 (42.7%) | 127 (35.4%) | 218 (38.1%) |
| Race/Ethnicity | Black or African American | 73 (34.3%) | 69 (19.2%) | 142 (24.8%) |
| Race/Ethnicity | Other | 49 (23.0%) | 162 (45.1%) | 211 (36.9%) |
| Race/Ethnicity | Unknown | 0 (0.0%) | 1 (0.3%) | 1 (0.2%) |
| Age at death (years) | Mean ± SD | 76.9 ± 11.9 | 79.0 ± 9.7 | 78.2 ± 10.6 |
| | Median [IQR] | 80.0 [70.0–86.0] | 81.0 [73.0–88.0] | 81.0 [72.0–87.0] |
| | Range | 43–90 | 50–90 | 43–90 |
| PMI (hours) | Mean ± SD | 11.5 ± 10.8 | 11.4 ± 18.4 | 11.4 ± 16.3 |
| | Median [IQR] | 7.0 [5.0–15.0] | 7.1 [4.5–13.9] | 7.0 [4.7–14.5] |
| Braak stage | I | 13 (6.1%) | 30 (8.4%) | 43 (7.5%) |
| Braak stage | II | 17 (8.0%) | 41 (11.4%) | 58 (10.1%) |
| Braak stage | III | 14 (6.6%) | 61 (17.0%) | 75 (13.1%) |
| Braak stage | IV | 17 (8.0%) | 32 (8.9%) | 49 (8.6%) |
| Braak stage | V | 50 (23.5%) | 45 (12.5%) | 95 (16.6%) |
| Braak stage | VI | 90 (42.3%) | 124 (34.5%) | 214 (37.4%) |
| | Missing | 12 (5.6%) | 26 (7.2%) | 38 (6.6%) |
| Thal phase | 1 | 2 (0.9%) | 13 (3.6%) | 15 (2.6%) |
| Thal phase | 2 | 12 (5.6%) | 9 (2.5%) | 21 (3.7%) |
| Thal phase | 3 | 11 (5.2%) | 15 (4.2%) | 26 (4.5%) |
| Thal phase | 4 | 21 (9.9%) | 14 (3.9%) | 35 (6.1%) |
| Thal phase | 5 | 110 (51.6%) | 64 (17.8%) | 174 (30.4%) |
| | Missing | 57 (26.8%) | 244 (68.0%) | 301 (52.6%) |
| TMT batch/plex | DLPFC | DLPFC: 24 plexes (b01–b24) | DLPFC: 24 plexes (b01–b24) | DLPFC: 24 plexes (b01–b24) |
| | STG | STG: 19 plexes (b01–b19) | — | — |

Notes: Values are n (%) for categorical variables, mean ± SD and median [IQR] for continuous variables.

Supplementary Table 2 is provided as an accompanying Excel workbook.
Supplementary Table 3 is provided as an accompanying Excel workbook.
Supplementary Table 4 is provided as an accompanying Excel workbook.
